# Genome-wide association study highlights escape from aphids by delayed growth in *Arabidopsis thaliana*

**DOI:** 10.1101/2022.11.10.515564

**Authors:** Chongmeng Xu, Yasuhiro Sato, Misako Yamazaki, Marcel Brasser, Matthew A. Barbour, Jordi Bascompte, Kentaro K. Shimizu

**Author notes:** These authors equally contributed to this study (Chongmeng Xu, Yasuhiro Sato). Co-correspondence; Y.S.; K.K.S.

## Abstract

Field studies have shown that plant phenological and architectural traits often explain substantial variation in herbivory. Although plant genes involved in physical and chemical defense are well studied, less is known about the genetic basis underlying effects of plant growth traits on herbivory. Here, we conducted a genome-wide association study (GWAS) of aphid abundance in a field population of *Arabidopsis thaliana*. This field GWAS detected a significant peak on the third chromosome of *A. thaliana*. Out of candidate genes near this significant genomic region, a mutant of a ribosomal gene (AT3G13882) exhibited slower growth and later flowering than a wild type under laboratory conditions. A no-choice assay with the turnip aphid, *Lipaphis erysimi*, found that aphids were unable to successfully establish on the mutant. These findings suggest the potential role of growth-related genes in altering herbivore abundance.

## Introduction

Plants are attacked by herbivores across their life cycle in natural environments. While chemical and physical traits have long been the main focus of anti-herbivore defense [1–4], plant life-history traits also account for herbivory variation in field environments [3,5,6]. For example, phenological changes (e.g., early flowering) can allow plants to escape from seasonal herbivory [7,8]. The visibility of plants for herbivores, namely plant apparency [9], also changes across plant ontogeny (e.g., from vegetative to reproductive phase), which alters the risk of herbivore attacks [6,10]. By focusing on genetic variation within a plant species, several studies have shown that plant phenological and architectural traits shape heritable variation in herbivory [4,11,12]. Yet, less is known about the underlying genetic basis of the plant life-history traits on herbivory.

Genome-wide association study (GWAS) is an effective way to dissect the genetic architecture of ecologically important traits [13,14]. Through associations between single nucleotide polymorphisms (SNPs) and traits, GWAS provides a hypothesis-free approach to identify novel genes from natural phenotypic variation [15,16]. Recent studies showed that controlled laboratory conditions are unlikely to reflect outdoor environments where interspecific interactions typically occur [17,18]. This fact emphasizes the importance of *in natura* study of functional genes [19–23]. To achieve this goal, it is important to conduct GWAS under field conditions.

*Arabidopsis thaliana* is a model plant species distributed and naturalized around the world. While *A. thaliana* usually blooms in spring after over-wintering, some cohorts have overlapping life cycles from spring to autumn [23–25]. When *A. thaliana* plants emerge from late spring to early summer, they are threatened by various herbivores [4,26]. Of the diverse insect herbivores, aphids are known to be major herbivores occurring across the natural distribution range of *A. thaliana* [27]. Because aphids often suck phloem sap from leaf veins and flowering stems, we hypothesized that plant life-history traits may play a key role in mediating aphid herbivory under field conditions.

To reveal the genetic architecture of aphid herbivory, we combined GWAS with a mutant analysis in *A. thaliana*. We first conducted GWAS of aphid abundance on 196 *A. thaliana* accessions grown in a field site in Zurich, Switzerland. To further test candidate genes, we then cultivated and released the turnip aphid *Lipaphis erysimi* on *A. thaliana* mutants.

## Materials & Methods

### Field GWAS

#### Plant genotypes

We obtained *Arabidopsis thaliana* genotypes that were selfed and maintained as inbred lines, called “accessions”, from the Arabidopsis Biological Resource Center (https://abrc.osu.edu/). We used the same set of 196 *A. thaliana* accessions as a previous study [28] except for two trichome mutants and an ungenotyped accession. All of these accessions were genotyped in the RegMap [29] and 1001 Genomes [30] projects. The list of accessions and phenotypes measured in this study is available in Table S1.

#### Field experiment

To observe aphids in a simulated late cohort, we exposed *A. thaliana* accessions to a field environment from 4 to 25 July 2018. This field experiment was conducted at Zurich, Switzerland, to use a field site within a native distribution range of *A. thaliana*. To keep all the accessions in the rosette stage at the start of the field experiment, we initially cultivated *A. thaliana* in a laboratory under a short-day condition (8h light/16h dark cycle at 20°C). Seeds were sown on 33-mm diameter Jiffy-seven^(R)^ pots and stratified under a constant dark 4°C condition for a week. Seedlings were cultivated in a growth chamber for 6 weeks under the short-day condition. Plants grown on the Jiffy-seven pots were then planted in a new plastic pot filled with mixed soils of agricultural composts (Profi Substrat Classic CL ED73, Einheitserde Co.) and perlites with compost to perlite ratio of 3:1 litter volume. Eight replicates of the 196 accessions were then transferred to the outdoor garden at the University of Zurich-Irchel (47° 23’N, 8° 33’E). Aphids were counted by a single observer every two or three days. To examine whether the aphid abundance differed between plants with and without flowering stems, we also recorded the presence or absence of bolting two weeks after the start of field experiment.

#### Data analysis

All GWAS analyses were performed using the GWA-portal (https://gwas.gmi.oeaw.ac.at) [31]. The target phenotype was the maximum number of aphids, which included *Lipaphis erysimi* and *Brevicoryne brassicae* (see Results), observed on a plant during the experiment. The imputed full-sequence dataset [31] was used as SNP data for the 196 accessions. Pseudo-heritability *h^2^* [31] was calculated for the target phenotype before association mapping. Accelerated mixed models [31] were used for association mapping with a correction of kinship structure. The number of aphids was ln(*x* + 1)-transformed to improve normality. After the association mapping, candidate genes were searched within ca. 5 kb near a focal SNP. To inspect organ-specific expression levels of candidate genes, we referred to Klepikova Arabidopsis Atlas [32] via The Arabidopsis Information Resource (https://www.arabidopsis.org/).

### Mutant analysis

#### *Arabidopsis thaliana* mutants

T-DNA sequence-indexed lines of *A. thaliana* were obtained from the Nottingham Arabidopsis Stock Centre (NASC) (https://arabidopsis.info/). In addition to Columbia-0 (Col-0, NASC Accession ID: N70000) wild type, we ordered four mutant lines for a ribosomal gene (AT3G13882) (Table S2). These original mutants were back-crossed to the Col-0 wild type three times. Following the instruction [33], we examined the insertion site by polymerase chain reaction (PCR) amplification and Sanger sequencing; and gene expression levels by Semi-quantitative reverse transcription and PCR (sqRT-PCR) (see captions of Fig. S1 and Fig. S2 for details). We found that one of the four lines, the SALK_039481 (NASC Accession ID: N670586), indeed had a T-DNA insertion on an exon of one of two splice variants (Fig. S1) and reduced expression level of AT3G13882 (Fig. S2), suggesting that the insertion disrupted the gene. In the other three lines, the insertion was not found or low germination rate prevented further experiments.

#### Laboratory experiments

To observe plant growth, we cultivated ten replicates of the ribosomal gene mutant and the Col-0 wild type under a long-day condition (16h light/8h dark cycle at 22°C/20°C). Seeds were sown on a 294 cm^3^ (= 7 × 7 × 6 cm) pot filled with the agricultural composts (Profi Substrat Classic CL ED73), and stratified at 4°C under a constant dark condition for a week. The stratified seeds were then transferred to the long day condition. Seedlings were grown for 20 days. The rosette diameter (cm) was recorded as an index of plant size before aphids were released, as described next.

To test whether aphids could establish a colony on the mutant plants, we released the turnip aphid *L. erysimi* on the wild type and the mutant of *A. thaliana*. The potted plants were separately enclosed with a mesh net. Five wingless females of adult aphids were released on each plant. The experimental aphids were derived from a source population established by a previous study [34]. The enclosed plants were incubated under the long-day condition. The number of aphids per plant was counted by eye 3, 7, 10, and 14 days after the release of aphids. We did not count aphids that escaped outside the area of a plant. Flowering time was defined as the number of days to flowering, and recorded during the aphid experiment.

#### Data analysis

We used linear mixed models (LMMs) or generalized linear mixed models (GLMMs) to test phenotypic differences between the mutant and the Col-0 wild type. The plant size and flowering time were analyzed using LMMs that assumed Gaussian errors. The number of aphids i.e., the count response was analyzed using GLMMs with Poisson error structure and log link function. Paired positions of a mutant and wild type plant were incorporated as a random effect to consider spatial heterogeneity within a growth chamber environment. An analysis of deviance with a F-test was used to test the significance of the mutant vs. wild type (df1 = 1) against phenotypic variation within the random effect of the paired positions (df2 = 10 pairs - 1 fixed effect = 9). All statistical analyses were performed using R version 4.0.3 [35]. For LMM and GLMM, we used the lmer and glmer function implemented in the lme4 package [36].

## Results

### Field GWAS of the aphid abundance

To monitor aphid abundance as well as visible plant traits, we transplanted 196 *A. thaliana* accessions in the field in Zurich within a native distribution range of *A. thaliana*. At the transplantation, all plants were at the rosette stage, i.e., no bolting occurred. After two weeks, 38% of individual plants initiated bolting, i.e., a stem was observed. The main herbivores were the two species of specialist aphids, *Lipaphis erysimi* and *Brevicoryne brassicae*. The aphid abundance was higher on bolted accessions than on non-bolted accessions (non-bolted and bolted plants = average 0.59 and 2.07 aphids, respectively; Welch’s *t*-test, *t* = –21.9, *df* = 941.2, *p* < 10^−15^), suggesting that plant life cycle might be associated with plants’ capacity to avoid aphids. In addition, we also distinguished the abundance of winged and wingless aphids to infer the colonization process of aphids on *A. thaliana*. Winged and wingless aphids were observed at the rosette stage at the first monitoring after the transplant, but many of these aphids did not establish a colony in subsequent monitoring (the days between 07 and 10 July 2018: Fig. S3). This observation suggests that colonized aphids do not always establish a colony and thereby the success of the colony establishment also depends on the presence of stem after colonization.

To reveal genetic architecture underlying variation in aphid abundance, we calculated heritability and then performed association mapping. Aphid abundance had high heritability among the plant accessions (h^2^ = 0.7), indicating the genetic control of this trait. Our mapping also detected a significant SNP in an intergenic region above the genome-wide Bonferroni threshold (chr3-4579292, *p* < 10^−8^, MAF=0.026: Fig. 1; see also Fig. S4 for quantile-quantile plots). Nearby this significant SNP (chr3-4579292), we found three candidate genes: such as a putative ribosomal gene (AT3G13882) that is homologous to a ribosome protein L34 gene (RPL34) [37], *EPIDERMAL PATTERNING FACTOR LIKE 3* (*EPFL3:* AT3G13898), and *MYB26*. Out of these three genes, the ribosomal gene (AT3G13882) is known to be highly expressed in vegetative organs such as leaves [32]. The other two genes, *EPFL3* and *MYB26*, are highly expressed only in reproductive organs such as anthers or pistils [32]. Because aphids were unlikely to suck saps from anthers and pistils, we focused on the ribosomal gene (AT3G13882) for further investigation.

**Figure 1.**
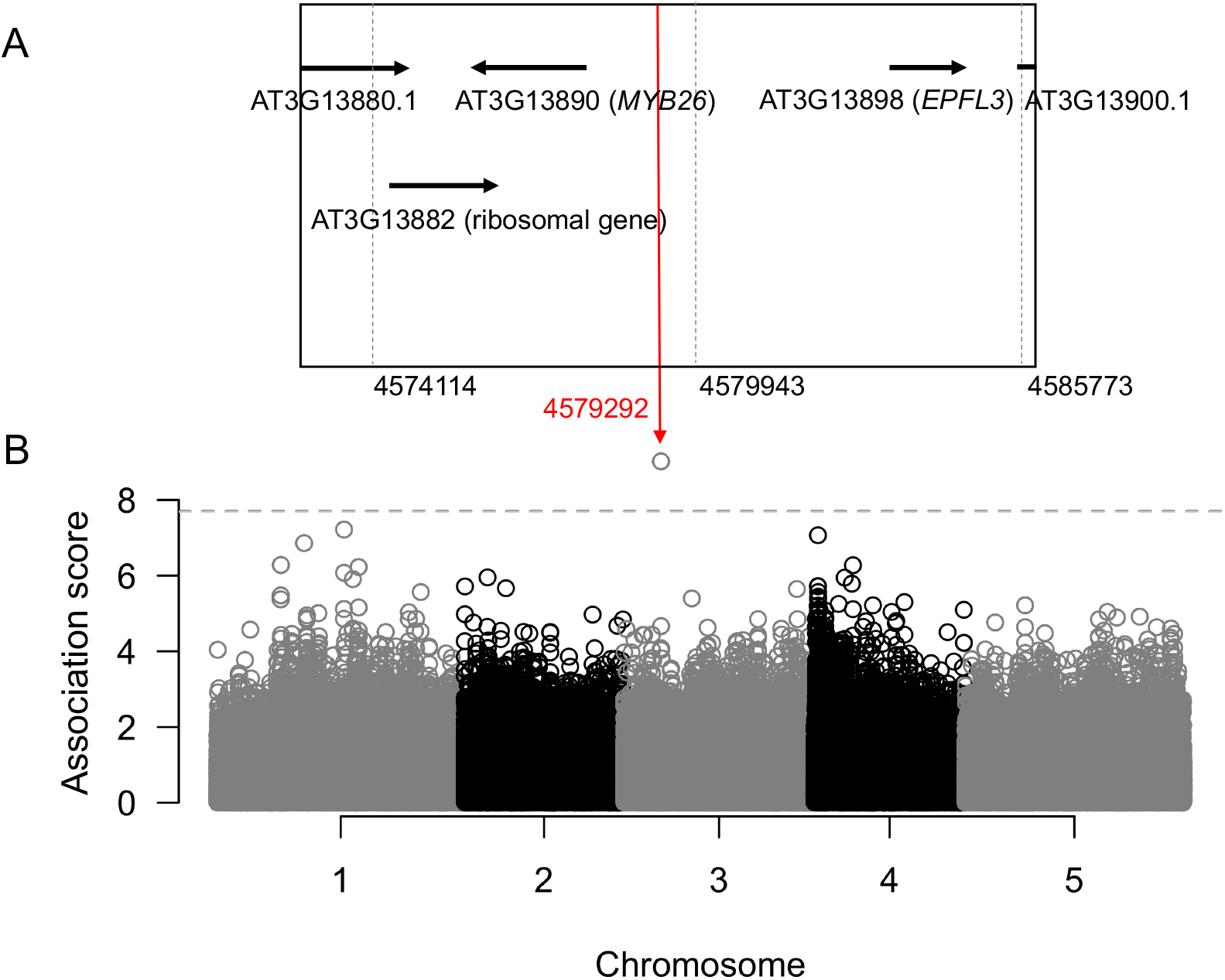
Genome-wide association study of aphid abundance on 196 Arabidopsis thaliana accessions grown in the field. (A) A genomic region within ca. 5 kbp from the top-scoring SNP at Chr3-4579292 displays the position of candidate genes. Only the longest splice variant (black horizontal arrow) is shown for each gene. (B) Manhattan plot shows the association score of-log_10_(p) across five chromosomes of A. thaliana with MAF cut-off at 0.025. A horizontal dashed line indicates the genome-wide Bonferroni threshold at p = 0.05.

### Mutant plant growth and aphid colony establishment in the laboratory

To examine visible phenotypes of the ribosomal gene mutant (AT3G13882), we compared growth and flowering time of this mutant with the Col-0 wild type. After 20 days of growth, the AT3G13882 mutant had a significantly smaller size than the wild type (*F_1,9_* = 42.1, *p* = 0.00011: Fig. 2A,B). The flowering time of the AT3G13882 mutant was also significantly later than the wild type (*F_1,9_* = 48.8, *p* < 0.0001: Fig. 2A,C). The slower growth and delayed flowering of the ribosomal gene mutant led us to test whether the delayed growth could prevent the establishment of aphid colonies after colonization.

**Figure 2.**
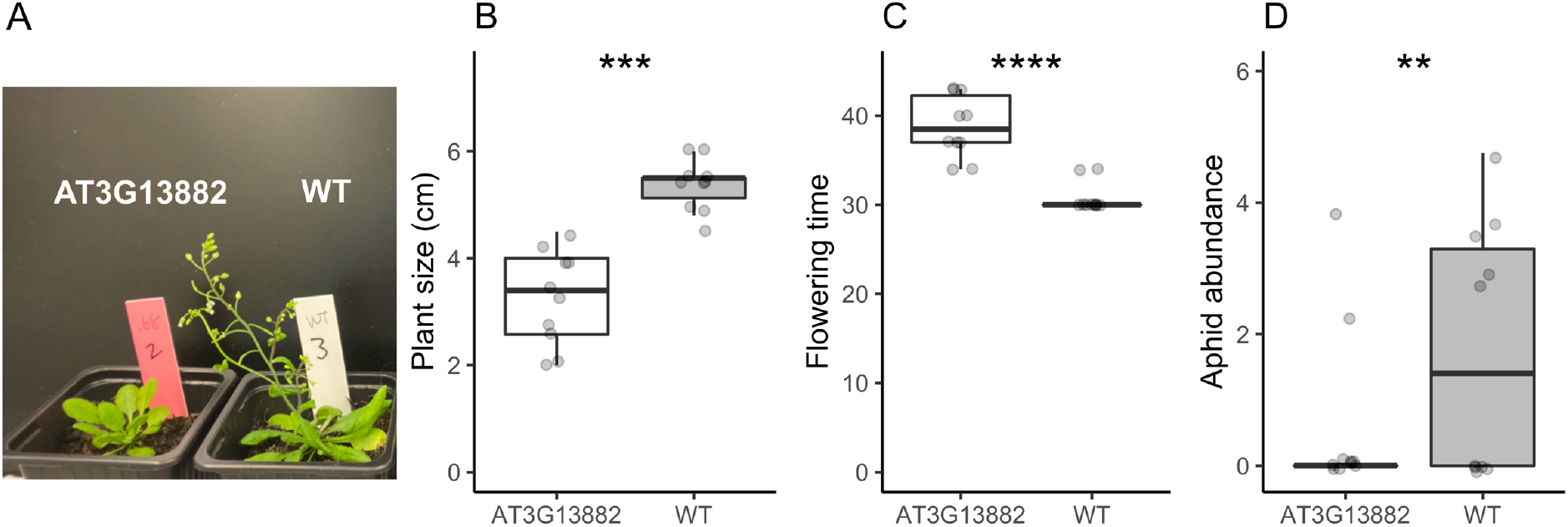
The photograph (A), plant size (B), flowering time (C), and aphid abundance (D) of the Col-0 wild type (WT) and the ribosomal gene mutant (AT3G13882) of Arabidopsis thaliana under laboratory conditions. Flowering time and aphid abundance represents the number of days to flowering and log_2_(no. of aphids + 1), respectively. Asterisks indicate the statistical significance by generalized linear mixed models; ** p < 0.01; *** p < 0.001; **** p < 0.001. Boxes: median with upper and lower quartile; Whiskers: 1.5 × inter-quartile range.

Then, to examine colony establishment after aphid colonization, we released wingless individuals of *Lipaphis erysimi* on rosette plants of the ribosomal gene mutant (AT3G13882) and the wild type. We observed a reduced number of aphids on the AT3G13882 mutant compared to the wild type at 7, 10, 14 days after the release of aphids (*F_1,9_* = 19.3, *p* = 0.0017 at 7 days; Fig. 2D: see also Fig. S5 for results at 10 and 14 days), suggesting that the delayed growth of the host negatively affected aphid colony establishment.

## Discussion

While ribosomal genes have long been considered housekeeping genes of the protein synthesis machinery, mutants of ribosome-related genes exhibit a wide variety of growth and reproductive phenotypes. For example, previous studies reported the reduction of leaf cell number [38], reduced root length [39], and the reduction of pollen number [16,40] regarding ribosomal gene mutations. In this context, our findings from the ribosomal gene AT3G13882 add insights into biotic interactions besides growth deterioration. We should note that further study on natural variants responsible for the delayed growth and reduced aphid abundance would be necessary to support its significance in the field. Multiple alleles on the AT3G13882 gene are also needed to provide strong evidence of those phenotypes. In the studies of a ribosomal gene *REDUCED POLLEN NUMBER1* (*RDP1*), null mutants showed a pleiotropic effect on plant growth and pollen number in *A. thaliana* [16,40]. Natural alleles of *RDP1* could alleviate pleiotropic growth defects [16]. In our study, the other growth-related genes or other mutations of AT3G13882 might have reduced the aphid abundance. Because transgenic approaches may not be effective to identify mutation sites affecting quantitative traits, further experimental tests, such as quantitative complementation [16,40], will be needed to identify natural causal variants that alter aphid abundance through delayed growth.

Overall, our study suggests the importance of plant life-history traits in altering herbivore abundance. Previous field studies used mutant plants to illustrate *in natura* roles of well-studied functional genes in chemical defense (e.g., *LOXs* in *Nicotiana attenuata* [2,3]) and physical defense (*GLABRA1* in *A. thaliana* [4]) against herbivores. In contrast, field GWAS offered a hypothesis-free approach to screen candidate genes responsible for herbivore abundance [14]. These lines of studies suggest that plant genetic variation governs herbivore abundance and communities [4,11,12,14], but a keystone gene shaping the ecological communities was not identified until recently [34]. Barbour et al. [34] have experimentally shown that pleiotropic effects of a glucosinolate biosynthesis gene *AOP2* on plant growth alter *A. thaliana*’s capacity to harbor aphids and their parasitoids. Our candidate genes might also exhibit pleiotropy on plant growth and secondary metabolism, but our findings support the notion that genes associated with plant growth can structure populations or communities of associated organisms. Since aphids and aphidophagous insects are widespread across terrestrial ecosystems [41,42], future studies may reveal cascading effects of delayed plant growth on food webs.

## Data availability

The data and codes are available at the GitHub repository (https://github.com/yassato/AraAphidGWAS).

## Authors’ contributions

C.X.: laboratory investigation, project administration, data curation, draft writing, review and editing; Y.S.: conceptualization, funding acquisition, project administration, supervision, field investigation, data curation, formal analysis, draft writing, review and editing; M.Y.: laboratory investigation, methodology (molecular), project administration, review and editing; M.B.: laboratory investigation, methodology (molecular); M.A.B: resources (insects), review and editing; J.B.: conceptualization, funding acquisition, supervision, review and editing; K.K.S.: conceptualization, funding acquisition, project administration, supervision, draft writing, review and editing.

## Acknowledgements

The authors thank L. Mohn, K.K. Thomsen, and all members of Shimizu group for their help with the establishment of field plots; and R. Hostettler for assistance with the molecular experiments.

## Supplementary Materials

**Figure S1.**
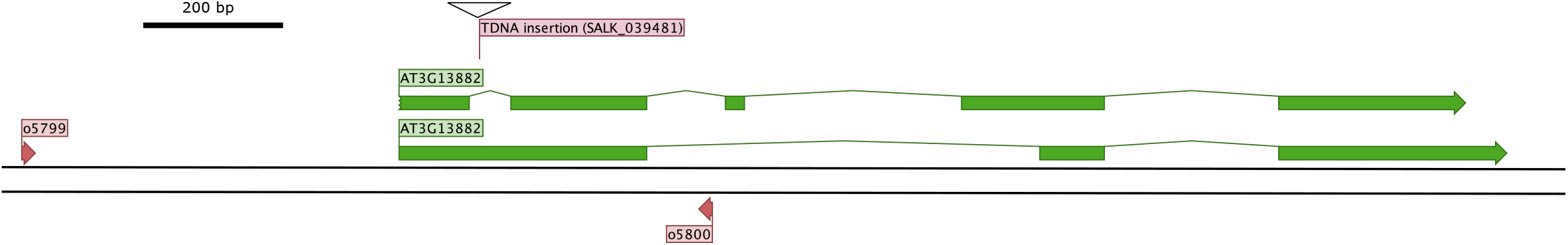
Schematic structure of the ribosomal gene AT3G13882. Green boxes: mRNA for two splice variants; triangle: the position of Transfer-DNA (T-DNA) insersion; red arrrows: primers for checking the T-DNA insertion. The primer IDs shown by four digits (oXXXX) correspond to those listed in Table S3. We downloaded the annotations from Genbank (Gene ID: 3768810), and visualized the position of the gene and primers using QIAGEN CLC Main Workbench. To confirm the T-DNA insertion site of SALK_O39481, we extracted DNA from leaves using the CTAB method. We then amplified the DNA by polymerase chain reaction (PCR) as follows: 2 min at 95° C; 35 cycles of 15 s at 95° C, 30 s at 55° C, 1.5 min at 72° C; and a final extension step of 3 min at 72° C. The PCR product was finally sequenced by Sanger sequencing to confirm the insertion site.

**Figure S2.**
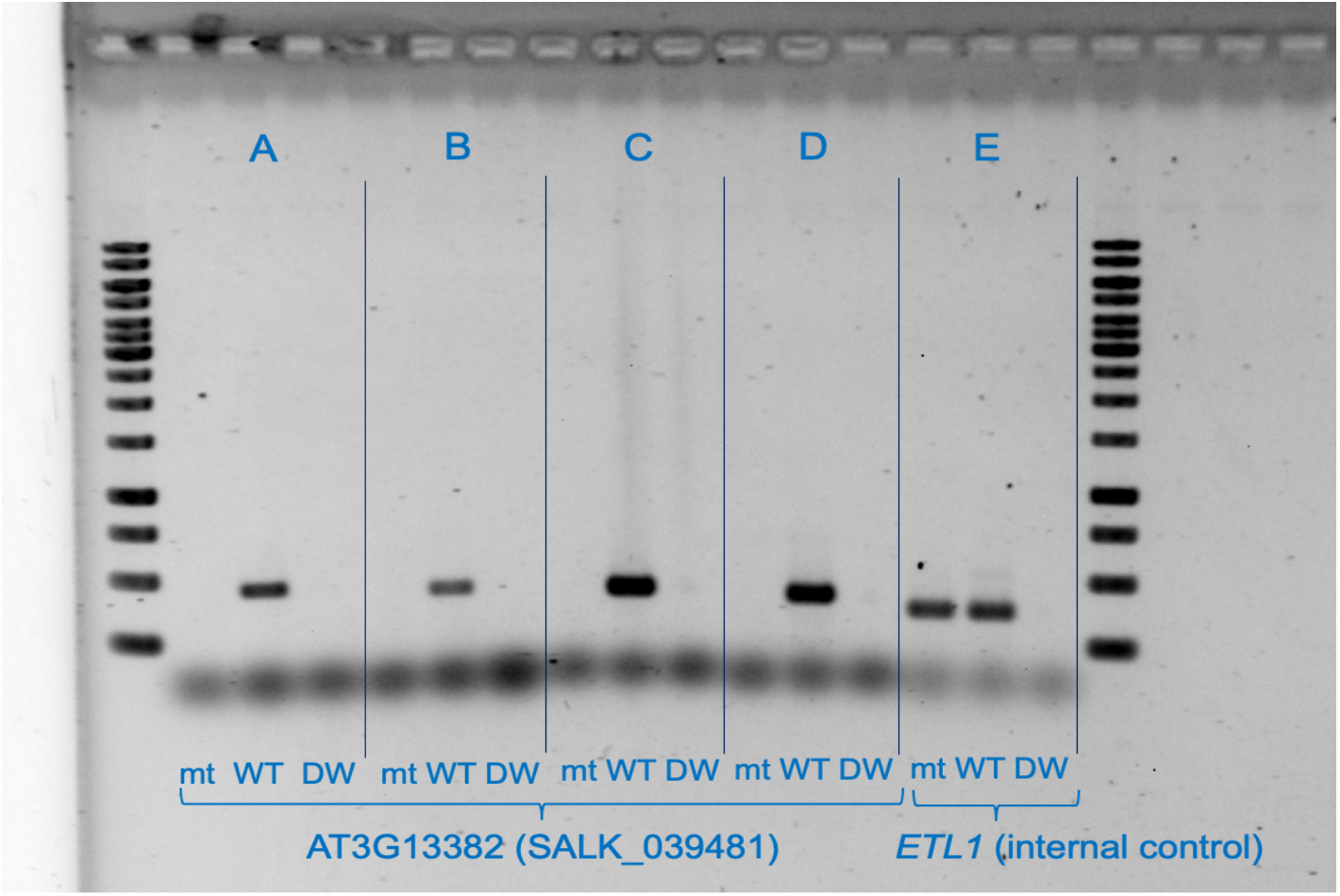
Semi-quantitative reverse transcription and polymerase chain reaction (sqRT-PCR) of the ribosomal gene mutant mutant AT3G13882 (SALK_039481). To perform sqRT-PCR, we extracted the RNA from leaves using RNeasy kit (Qiagen: Catalogue Number: 74181) and purified the RNA with DNA-free kit (Ambion: Cat. No. AM1906). RNA concentration was measured by Qubit spectrophotometer (Invitrogen: Cat. No. Q10211). Then we obtained the cDNA using High-Capacity RNA-to-cDNA kit (Applied Biosystems: Cat. No. 4387406) from 500 ng of the total RNA. The cDNA was amplified by PCR as follows: 3 min at 95° C; 28 cycles of 15 s at 95° C, 30 s at 55° C, 1 min at 72° C; and a final extension step of 5 min at 72°C. A gel electrophoresis was performed using a 1% agarose with 120 V for 60 min. The PCR products were finally visualized an UV trans-illuminator system. Two primer sets A-D were used for AT3G13882; and E for an internal control gene ETL1 (see Table S4 for the primer information). This figure shows that the gene expression of AT3G13882 is suppressed in the mutant line SALK_039481.

**Figure S3.**
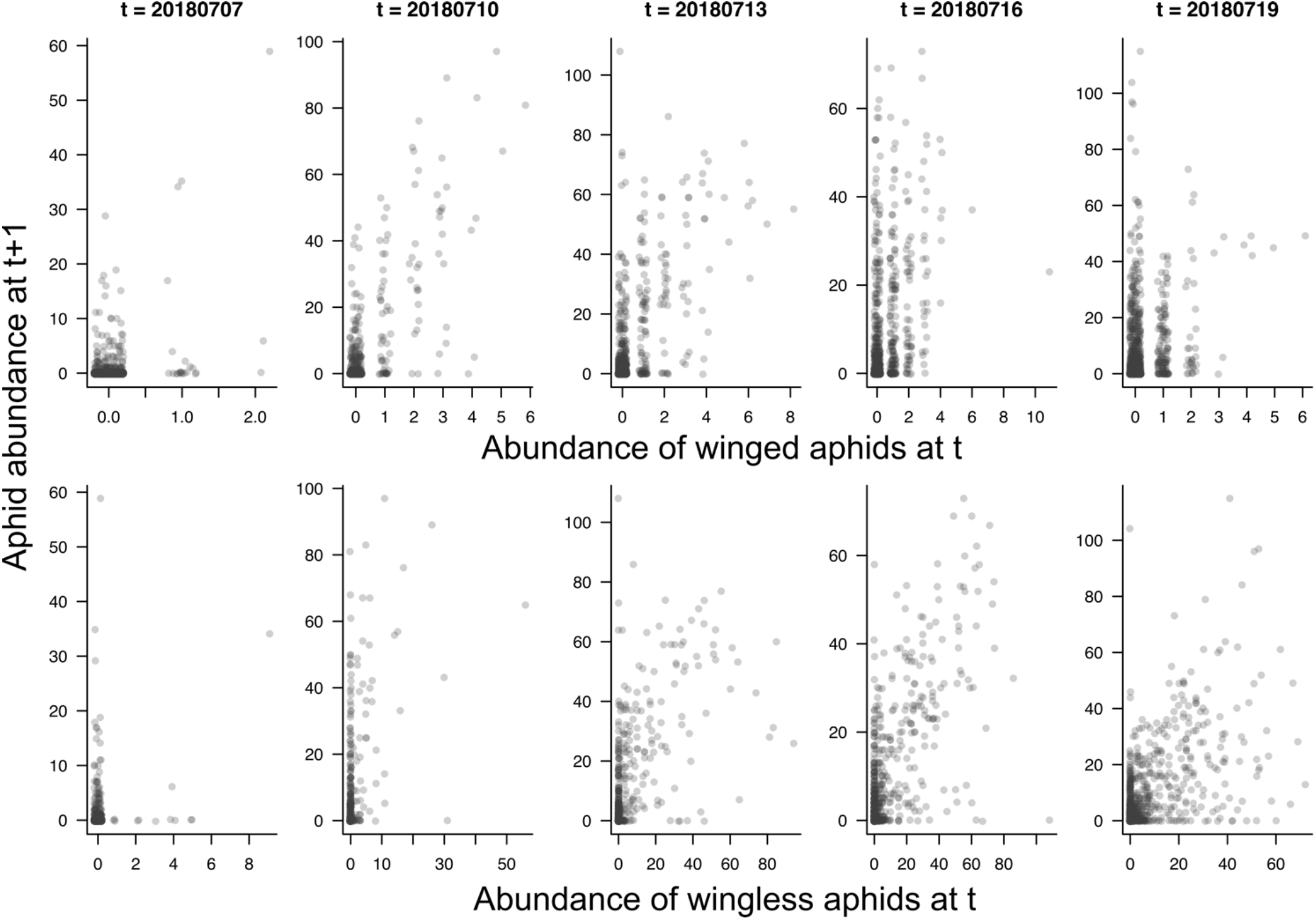
Temporal patterns of the emergence of wingless and winged aphids during the field GWAS experiment. X-axes show the number of winged (top row) or wingless (bottom row) aphids at current monitoring day (t=year-month-day). Y-axes show the total number of both winged and wingless aphids at the next monitoring date (t+1). A single point corresponds to an individual plant. The number of aphids represents the total number of individuals of both *Lipaphis erysimi* and *Brevicoryne brassicae*.

**Figure S4.**
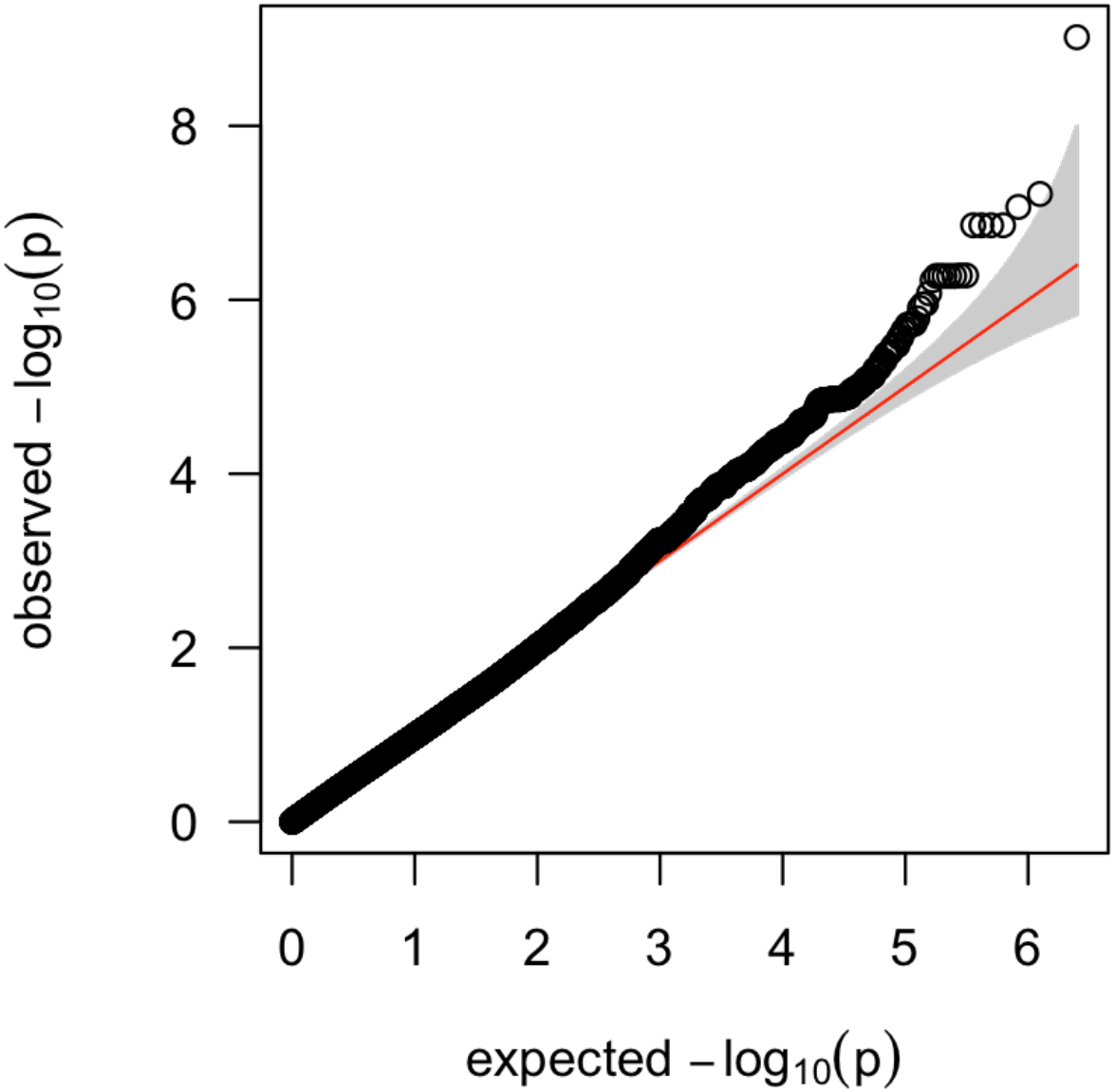
A quantile-quantile (QQ) plot showing relationships between the observed and expected association score of -log_10_(p). A red solid line indicates randomly expected association scores abd the shaded area corresponds to its 95% confidence intervals. This figure shows that the top-scoring SNP at Chr3-4579292 is larger than the upper 95% confidence interval.

**Figure S5.**
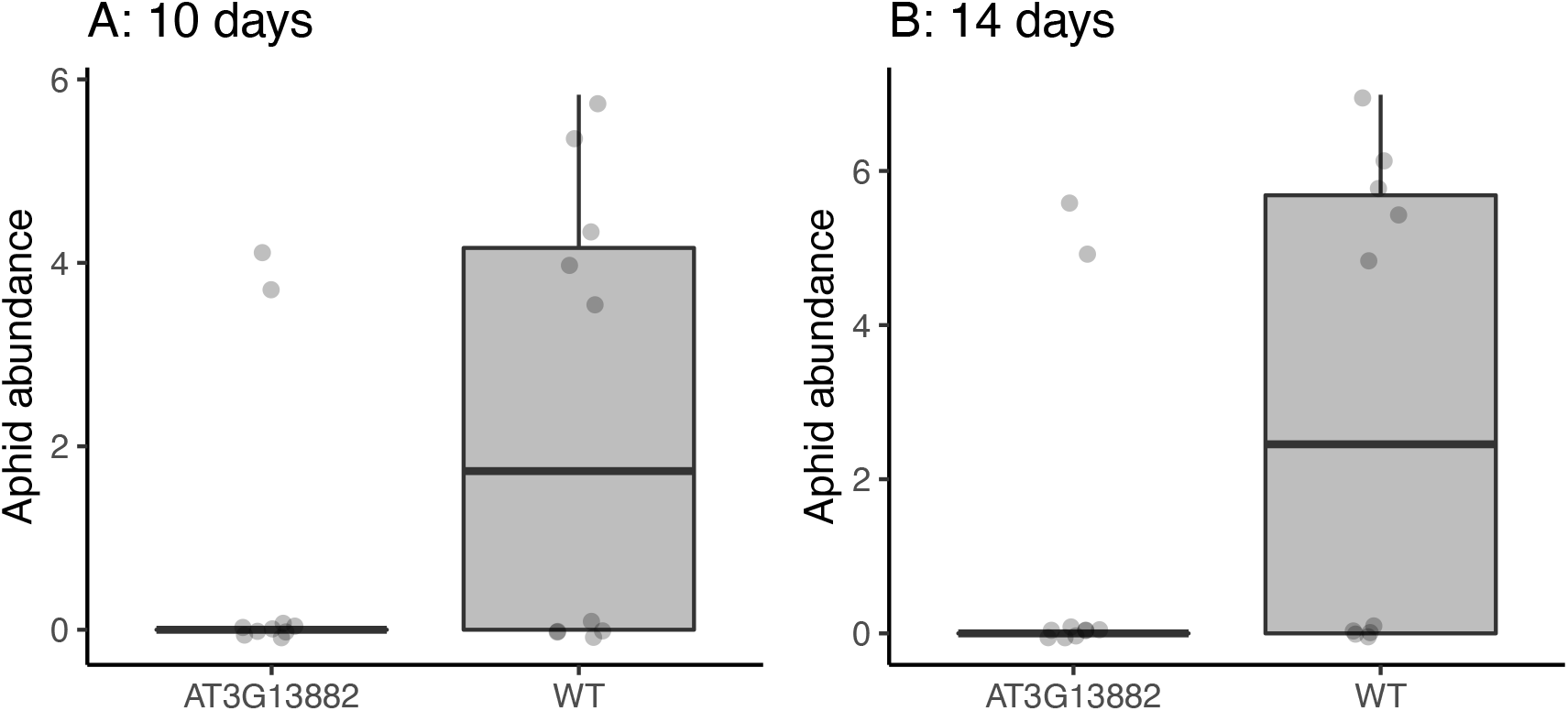
Aphid abundance during the later period of incubation. Aphid abundance represents log_2_(no. of aphids + 1). Same as the result at 7 days (Fig. 2D), both the left and right panels have significant differences of the aphid abundance between the mutant (AT3G13882) and wild type (WT) (F_1,9_ = 56.96, p < 10^−4^ for 10 days; F_1,9_ = 131.3, p < 10^−5^ for 14 days).

Table S1. List of GWAS accessions and phenotypes.

Table S2. List of mutant lines examined in this study.

Table S3. List of primers to check the T-DNA insertion or gene expression.

Table S4. Primer set information for sqRT-PCR results shown in Figure S2.

